# Decode-gLM: Tools to Interpret, Audit, and Steer Genomic Language Models

**DOI:** 10.1101/2025.10.31.685860

**Authors:** Aaron Maiwald, Piotr Jedryszek, Florent Draye, Bernhard Schölkopf, Garrett M. Morris, Oliver M. Crook

## Abstract

While genomic language models are enabling the *de novo* design of entire genomes, they remain challenging to interpret, limiting their trustworthiness. Here, we show that sparse autoencoders (SAEs) trained on Nucleotide Transformer activations decompose hidden representations into interpretable biological features without supervision. Across layers and model sizes, SAEs identified over 60 diverse functional annotations encoded in the model’s activations. This included viral regulatory elements such as the CMV enhancer, despite viral genomes being excluded from training data. Tracing this signal revealed contamination in reference databases, demonstrating that interpretability methods can audit training data and identify hidden data leakage. We then show that Meta-SAEs, trained on the decoder weights of another SAE, can identify conceptual hierarchies encoded in the model, including a more abstract feature related to multiple HIV annotations. We confirmed that the features identified by our SAEs were learned during pretraining through probing a randomly initialised model. Finally, we demonstrate that our SAEs allow us to steer model predictions in biologically meaningful ways, showing that we can use an antibiotic-resistance SAE-feature to steer the model toward the A1408G aminoglycoside-resistance mutation in the ribosomal gene 16S rRNA. Together, these results establish SAEs as a method for both discovery and auditing, providing a toolkit for interpretable and trustworthy genomic foundation models. Readers can explore our findings at https://interpretglm.netlify.app/.

## 1 Introduction

For the first time, we can intelligently compose entire, functional bacteriophage genomes using artificial intelligence (King et al., 2025). This achievement is the culmination of rapid advances in improving genomic language models (Benegas et al., 2023; Brixi et al., 2025; Dalla-Torre et al., 2025), training them to predict masked nucleotides across thousands of genomes. This progress mirrors breakthroughs using protein language models (Ferruz et al., 2022; Lin et al., 2023; Nijkamp et al., 2023), capable of designing novel, functional fluorescent proteins (Hayes et al., 2025) or lysozymes (Madani et al., 2023).

During pretraining, genomic language models learn to encode high-level genomic functions in their embeddings (Nguyen et al., 2023; Benegas et al., 2023; Brixi et al., 2025; Dalla-Torre et al., 2025). Given that a large fraction of even the human genome remains uncharacterised, genomic language models may encode functional information beyond our current scientific understanding. Decoding what genomic language models have learned could then be a new route to scientific insight in genomics. One approach to understanding neural network representations is *probing* —training classifiers to predict specific input properties from model representations (Alain & Bengio, 2018; Immer et al., 2022). High predictive performance of such classifiers is taken to suggest that the model encodes the corresponding concept. While effective for testing whether representations encode known concepts, probing requires labelled data. For uncharacterised genomic regions, such functional labels are, by definition, absent, making supervised methods less useful.

Sparse autoencoders (SAEs) offer an unsupervised approach to interpreting model representations by learning a sparse linear decomposition of internal activations, in which each input is expressed using only a small number of latent directions. This approach has been highly successful for text-based language models(Yun et al., 2023; Bricken et al., 2023; Cunningham et al., 2023). Unlike standard autoencoders that compress inputs, SAEs project them into higher-dimensional spaces to overcome *polysemanticity* —the fact that, in neural networks, individual neurons usually activate on multiple, unrelated input features. Elhage et al. (2022) hypothesise that polysemanticity allows the model to encode more features than it has neurons, by distributing the representation of features. By expanding the representational space, SAEs disentangle these features, allowing individual dimensions of that space (*SAE-latents*) to capture individual input features more faithfully than the original neurons, thereby increasing interpretability (see Figure 1a and Related Work A for details).

**Figure 1:**
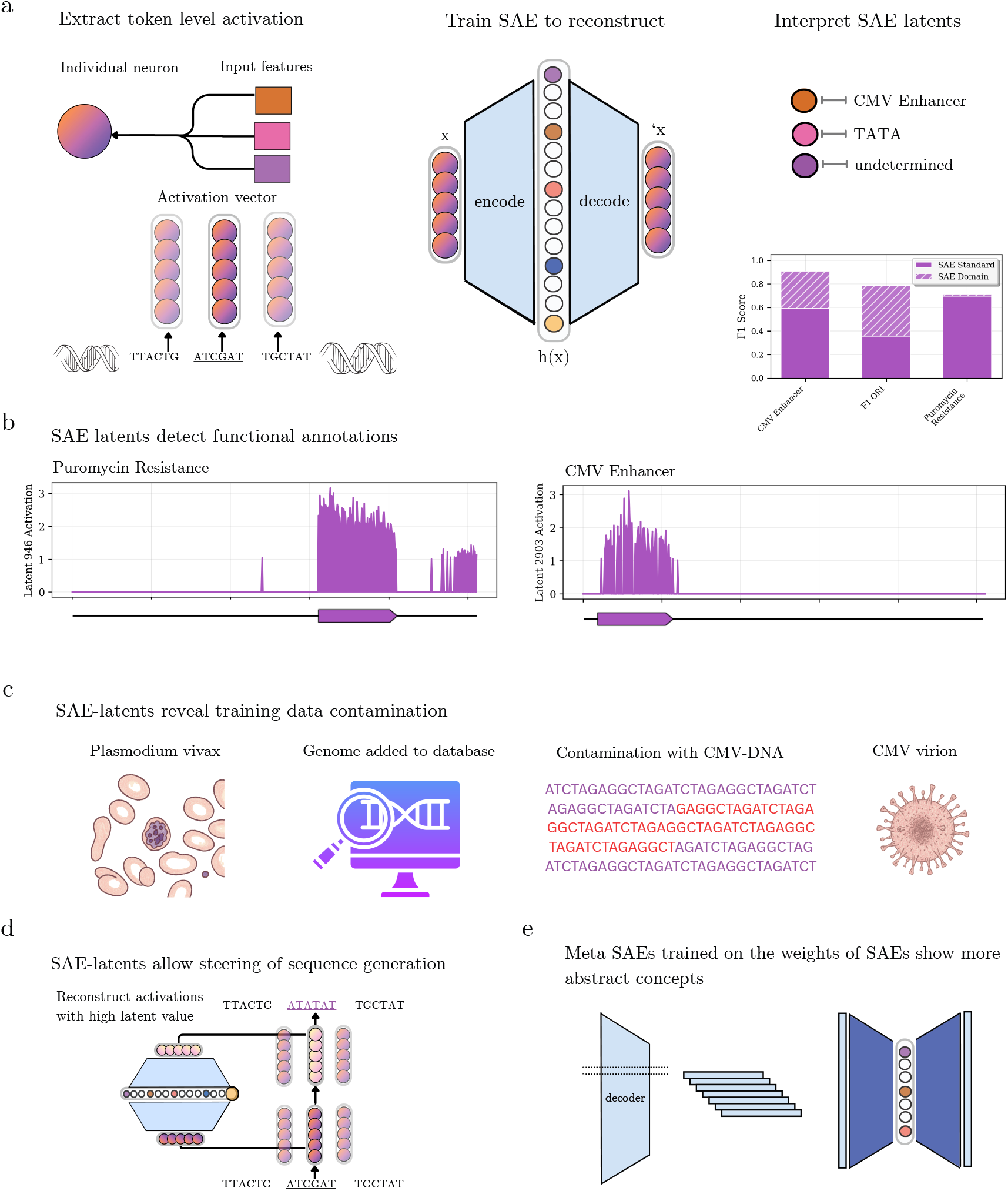
Interpreting Genomic Language Models using Sparse Autoencoders. a) Left: Individual neurons tend to be hard to interpret as they activate on multiple unrelated input features. Middle: Sparse autoencoders (SAEs) can project activation vectors into a higher-dimensional, sparse space, in which individual dimensions (“latents”) associate more closely with single input features. Right: In some cases, these latents correspond to known biological concepts, though not always. b) Two examples of latents activating selectively on puromycin resistance genes (left) and CMV enhancers (right). c) The model has learned to encode CMV enhancers, despite viral genomes being intentionally excluded from the training data. We trace this back to contamination of the training data with viral DNA. d) We show that our SAEs can be used to steer model generation by intervening on model activations. e) Meta-SAEs are trained on the decoder weights of other SAEs and sometimes decompose them into more abstract features.

This approach has been highly productive in the domain of protein language models. Simon & Zou (2025) and Gujral et al. (2025) apply this technique to ESM-2, uncovering thousands of encoded features ranging from entire protein families like the PTH family to biophysical properties like zinc finger domains. Comparable analyses of genomic language models remain limited. Contemporaneous with this work, Brixi et al. (2025) applied sparse autoencoders to the genomic model EVO-2, identifying SAE-latents moderately to strongly associated with frameshifts, coding sequences, introns, and exon starts and exon ends, among others.

We make four main contributions. First, we demonstrate that SAEs can discover an extensive vocabulary of biological concepts from the activations of the Nucleotide Transformer (Dalla-Torre et al., 2025), far larger and more granular than previously shown for genomic language models. In particular, SAEs identify over 60 biological concepts with F1-scores above 20%. Second, we show that this includes unexpected viral elements that we trace back to training data contamination. Third, we show that Meta-SAEs, trained to sparsely encode and reconstruct SAE decoder weights, can identify a more abstract concept related to HIV. Fourth, we verify the causal role of these features within the Nucleotide Transformer through causal interventions. In particular, we demonstrate that some SAE-features related to antibiotic-resistance enable biologically meaningful forms of steering, inducing the model to shift probability toward the A1408G aminoglycoside-resistance mutation in the ribosomal gene 16S rRNA. Finally, we provide a web interface for readers to explore the features we discovered at https://interpretglm.netlify.app/.

## 2 Results & Discussion

We analysed the Nucleotide Transformer (Dalla-Torre et al., 2025), a bidirectional encoder model pretrained on 850 genomes spanning multiple kingdoms of life. The model achieves strong performance across a range of tasks, indicating that it has captured extensive genomic structure. Prior work showed that its embeddings cluster into a small number of broad functional categories, including introns and coding sequences, but given the diversity of its training data, this likely represents only a fraction of the biological concepts it encodes.

Sparse autoencoders (SAEs) provide a method for disentangling such representations. Trained under a sparsity constraint so that only a small subset of neurons activate for any given input, SAEs typically project inputs into a higher-dimensional space, allowing features distributed across multiple neurons to be separated into components associated with a single input feature. In the natural language domain, this approach identified SAE-latents activating on features ranging from sycophancy to the Golden Gate Bridge (Templeton et al., 2024). Here, we adapt this approach to genomic language models.

To train SAEs, we extracted activations from the Nucleotide Transformer using plasmid sequences from AddGene (Kamens, 2015). Plasmids are absent from Nucleotide Transformer’s pretraining data, creating a distributional shift, and can be densely annotated with functional elements that are useful to characterise SAE-latents. Following previous work, we chose to extract multi-layer perceptron (MLP) outputs. For a single input of *n* tokens, these consist of *n* vectors of dimension *d*_*mlp*_, where *d*_*mlp*_ is the dimensionality of the MLP outputs. To monitor how features change throughout the model, we extracted activations across transformer blocks. We then trained SAEs to reconstruct MLP outputs while maintaining sparse activation, thereby learning an overcomplete dictionary of latent features. After hyperparameter tuning, we selected the SAEs with the best combination of reconstruction and sparsity loss.

To interpret the SAE-latents, we examined their most strongly activating tokens on two non-overlapping test sets (1,000 plasmids each) and associated them with functional annotations from pLannotate (McGuffie & Barrick, 2021). We then quantified the strength of the association between SAE-latents and annotations using F1-scores. For more details, see Methods B.

### 2.1 Discovering Unexpected Concepts using Sparse Autoencoders

To efficiently evaluate thousands of SAE latents for their association with annotations, we used the following heuristic. Our pipeline examined the 20 most activating tokens for each SAE latent and flagged any latent where at least half of the most activating tokens share an annotation. For each flagged latent, we then compute the F1-score of the latent-annotation pair, selecting an activation threshold on the validation set that maximises F1-score and then measuring F1-scores on separate test sets.

Based on previous studies, we expect feature representations to change across layers. We therefore train SAEs across the first, fourth, seventh, tenth and twelfth (final) MLP-layer^1^ of the smallest Nucleotide Transformer (50m). We call an SAE-latent interpretable if it has an F1-score of at least 20% for some annotation. We find that the number of interpretable SAE-latents varies strongly across layers, with none in the first layer and peaking in the 7th layer with over 60 (see Fig. 2b). This is consistent with previous findings that middle-to-late layers encode the most extractable features (Simon & Zou, 2025). Across layers, the number of interpretable SAE-latents far outnumbers the number of interpretable MLP-latents, and the most interpretable SAE-latents have consistently far higher F1-scores (see Fig. 2a). This confirms that our SAEs increase interpretability compared to the base model MLP.

**Figure 2:**
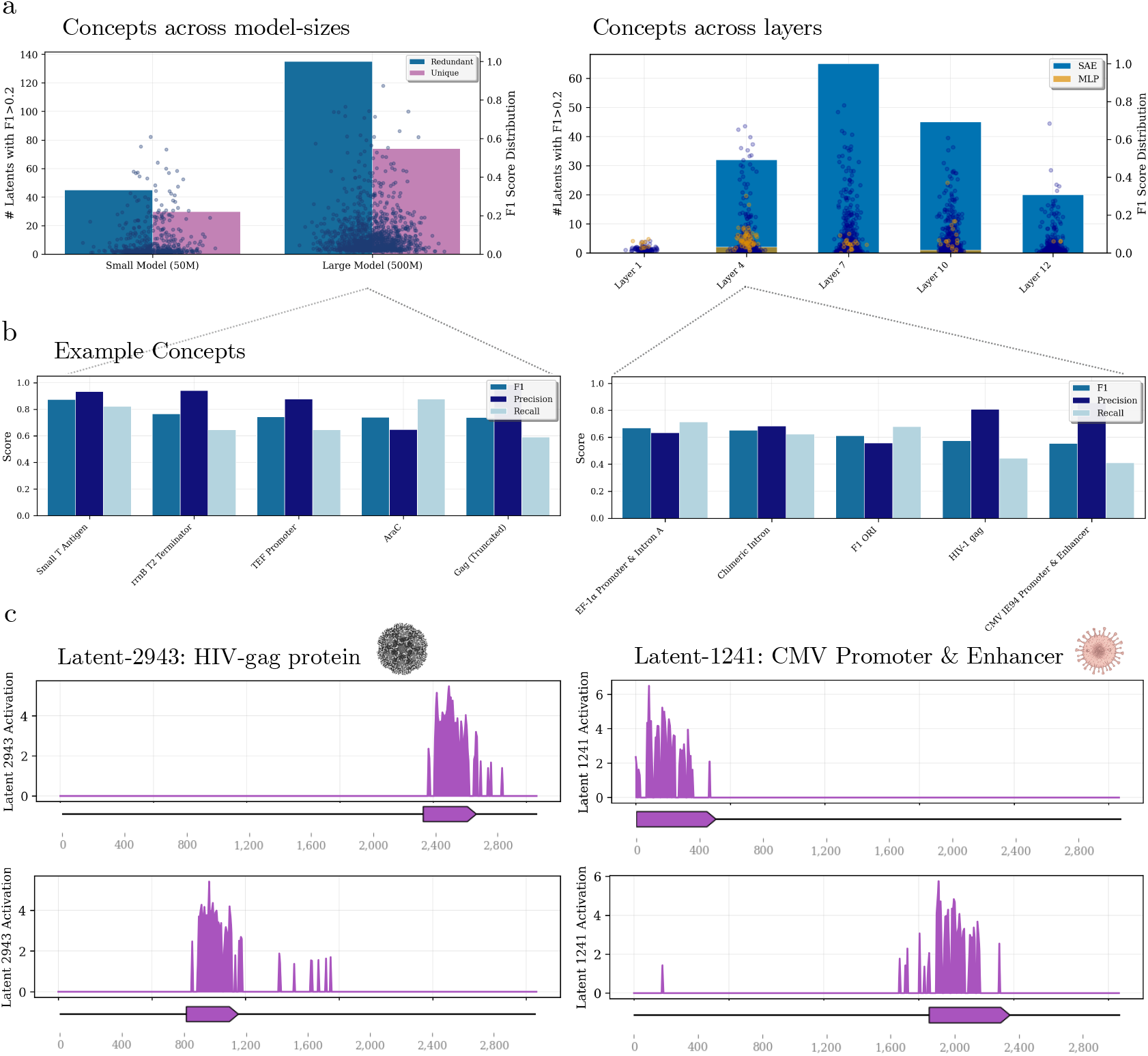
Sparse Autoencoders extract a large set of interpretable biological concepts across layers and model sizes. **a)** Left: we compare the number of interpretable SAE-latents across two model sizes (50m vs 500m) at the same model location (layer 10), as well as the distribution of F1-scores across latents. Right: shows how the number of interpretable latents and the distribution of F1-scores change across layers of the small model. As a baseline, we compare to the number of interpretable MLP-latents (yellow). **b)** Shows the five SAE-latents with the highest F1-scores in the large Nucleotide Transformer (left) and the fourth layer of the small model (right). **c)** Shows examples of two SAE-latents firing selectively on genes coding for the HIV-gag protein (left) and on CMV Promoters and Enhancers (right).

We expected a larger Nucleotide Transformer to encode more input features. We therefore trained SAEs on the 10th layer of the 500m Nucleotide Transformer, using the same hyperparameters as before. As expected, we find that our SAEs extract far more interpretable latents from the larger model. To save computational cost, we only train an SAE on layer 10 of this larger model. We find 135 unique latent-concept pairs, versus at most 65 in any layer of the smaller model. Secondly, the most interpretable SAE-latents have a much higher F1-score for the large model than for the smaller model: 87.3 % (for the small T-antigen) versus 77.9 % (for Puromycin Resistance). For comparison, a latent that fires independently of an annotation appearing in the input would have an expected F1 score below 2.0% for the annotations in our dataset.^2^

Unexpectedly, some of the SAE-latents with the highest F1 score were associated with viral annotations—CMV enhancer (F1: 59%) and HIV-gag with 57.3% F1-score. The CMV enhancer, ubiquitous in mammalian expression vectors, drives strong gene expression for research and gene therapy applications. HIV Gag proteins are structural polyproteins that assemble to form the core components of new HIV virus particles, playing a central role in viral assembly, budding, and maturation. These features appear despite Dalla-Torre et al. (2025) explicitly excluding viral genomes from training. To confirm that these features are encoded via pretraining, we therefore tested how well they can be extracted from the trained Nucleotide Transformer compared to a randomly initialised model of the same architecture. To do this, we train logistic regressions on the Nucleotide Transformer’s MLP outputs (“linear probes”) for both conditions. For each concept shown in 2b, we find that across 30 probes in each condition, the average F1-score is far greater for the pretrained model. The smallest increase in F1-score is an absolute 17.2 %, and the average increase is over 50 % (see Supp. Figure 5 for details.) This confirmed that these features are encoded via pretraining.

We consider two explanations for how the Nucleotide Transformer could have learned to encode CMV features during pretraining despite viral exclusion: (1) the model has somehow extrapolated from non-CMV-related DNA, (2) database contamination introduced CMV DNA into supposedly virus-free reference genomes. We next examine the second explanation.

### 2.2 Explaining the Source of Viral Features

To determine whether CMV enhancer sequences appeared in the training data despite viral genome exclusion, we used BLAST to query all CMV enhancer sequences from our plasmid dataset against the model’s training corpus. We identified five entries with near-perfect matches to the CMV enhancer (Table 1, each with E-values indicating extreme statistical significance ≤ 2.1 *×* 10^−15^). The matching organisms—bacteria and one protozoan—cannot be naturally infected by CMV, which exclusively targets mammalian cells. This makes viral integration unlikely and points to database contamination as the source. Such contamination occurs through multiple pathways: during sample collection (from infected organisms), laboratory preparation (via shared equipment or cross-contamination), or sequence assembly. This problem affects approximately 0.54% of GenBank entries (Steinegger & Salzberg, 2020), suggesting a 98.9% probability that at least one of the 850 training genomes contains contamination.

**Table 1:**
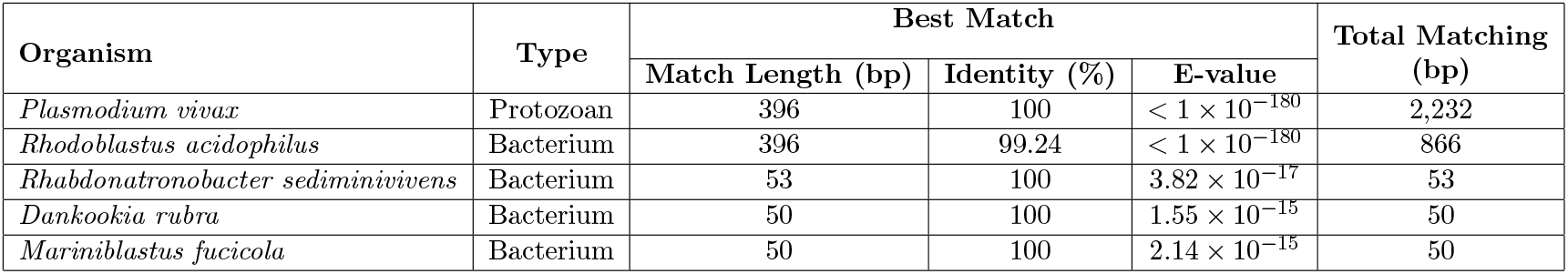
CMV Enhancer Matches in Training Data. BLAST Match Statistics from querying all CMV enhancer sequences in our test set against the Nucleotide Transformer’s training dataset, finding five organisms with significant matches.

**Table 2:** SAE Evaluation Metrics Across Layers.

| Model Size | Layer | NMSE | L1 Loss | Dead Latents (%) |
| --- | --- | --- | --- | --- |
| 50M | L1 | 0.0075 | 0.0293 | 5.74 |
| 50M | L4 | 0.0842 | 0.0168 | 43.43 |
| 50M | L7 | 0.0682 | 0.0396 | 36.13 |
| 50M | L10 | 0.0846 | 0.0528 | 62.72 |
| 50M | L12 | 0.0743 | 0.1045 | 50.10 |
| 500M | L10 | 0.1179 | 0.0213 | 77.72 |

Our discovery of training data contamination demonstrates three critical points. First, interpretability tools such as SAEs can serve as auditing mechanisms, allowing researchers to identify and remove hidden sequences that may compromise model behaviour and downstream analyses. Second, this finding highlights a biosecurity concern: even when viral genomes are explicitly excluded, viral information can leak back into training corpora through contaminated reference sequences, undermining safeguards that rely on dataset curation alone. Third, contamination complicates scientific claims about generalisation, since models may appear to succeed on supposedly novel test organisms that were, in fact, inadvertently represented during training. Together, these points emphasise both the promise of interpretability for ensuring data integrity and the need for more rigorous standards in constructing biological training datasets.

While SAEs may help to surface data contamination, directly scanning the data for the suspected contamination using a sequence alignment-based approach is probably faster and more sensitive. However, there may be scenarios in which model developers withhold training data, for example, because some of it is proprietary or subject to privacy constraints. In such cases, it is not possible to scan the training data, and here SAEs may be useful to detect issues with training data composition.

The identified CMV enhancer contamination represents only 500 of 29 billion training tokens— perhaps insufficient to fully account for the model’s strong CMV enhancer representation (78% F1-score). The remaining signal likely stems from either additional undetected contamination or transfer learning from similar mammalian enhancer sequences. To better understand the nature of this learned feature, we characterised the sequence properties that the CMV enhancer probe relies on. To do this, we ablated known regulatory motifs and compared the decrease in probe probability to random ablation. Our analysis revealed that the probe’s performance does not primarily rely on known regulatory motifs (see Supp. Material C). To test if the probe instead relied on local k-mer statistics, we used u-shuffle (Jiang et al., 2008) to create decoy sequences, preserving k-mer frequencies while disrupting their order. We found that preserving 5-mer frequencies was sufficient to recover 85% of the original probe probability, suggesting that the model represents the enhancer through k-mer sequence patterns rather than biologically known motifs (see Supplementary Material C for the full ablation and decoy analysis).

### 2.3 Examining Conceptual Hierarchies with Meta-SAEs

Building on SAEs, Meta-SAEs (Leask et al., 2025) are trained on the decoder-column weights of another SAE, which are intended to encode input-features. The Meta-SAE encoder, thereby, is designed to decompose features into more abstract, constituent features. To apply Meta-SAEs to our single-annotation features, we train Meta-SAEs on our SAE for the large Nucleotide Transformer (500m), training exclusively on decoder-columns of SAE-latents with positive F1 scores, sweeping across expansion factors and sparsity coefficients. After comparing multiple methods, we find that JumpReLU (*d*_*meta*−*sae*_ = 128) achieves the best trade-off between L0 sparsity (3-6) and explained variance (30–40%). See Methods B for details. To interpret the Meta-SAE latents, we inspect the top-20 most activating “child-latents” for each meta-latent.

We find that several Meta-SAE latents activate on clusters of semantically related child-latents. One striking example is latent #107, which activates most strongly on child-latents associated with HIV-related annotations: the Rev Response Element (RRE), HIV-1 dimerisation initiation sequence (HIV-1 DIS), HIV-1 psi packaging element (HIV-1 psi), HIV primer binding site (HIV PBS), and envelope proteins.

To measure the association between latent #107 and this set of annotations, we extracted the corresponding encoder row and evaluated it as a linear classifier. To do this, we first determined an optimal bias on a validation set of annotated sequences and then measured the F1-score on a hold-out test set of 500 annotated plasmid sequences. As a control, we also measured the F1-score for every subset of annotations, to assess if the latent corresponds to a more specific set of features. As shown in Fig. 3, we find that the latent’s F1-score is relatively high for all the annotations (57.2%) and substantially higher than for any individual annotation (≤ 44.8%).

**Figure 3:**
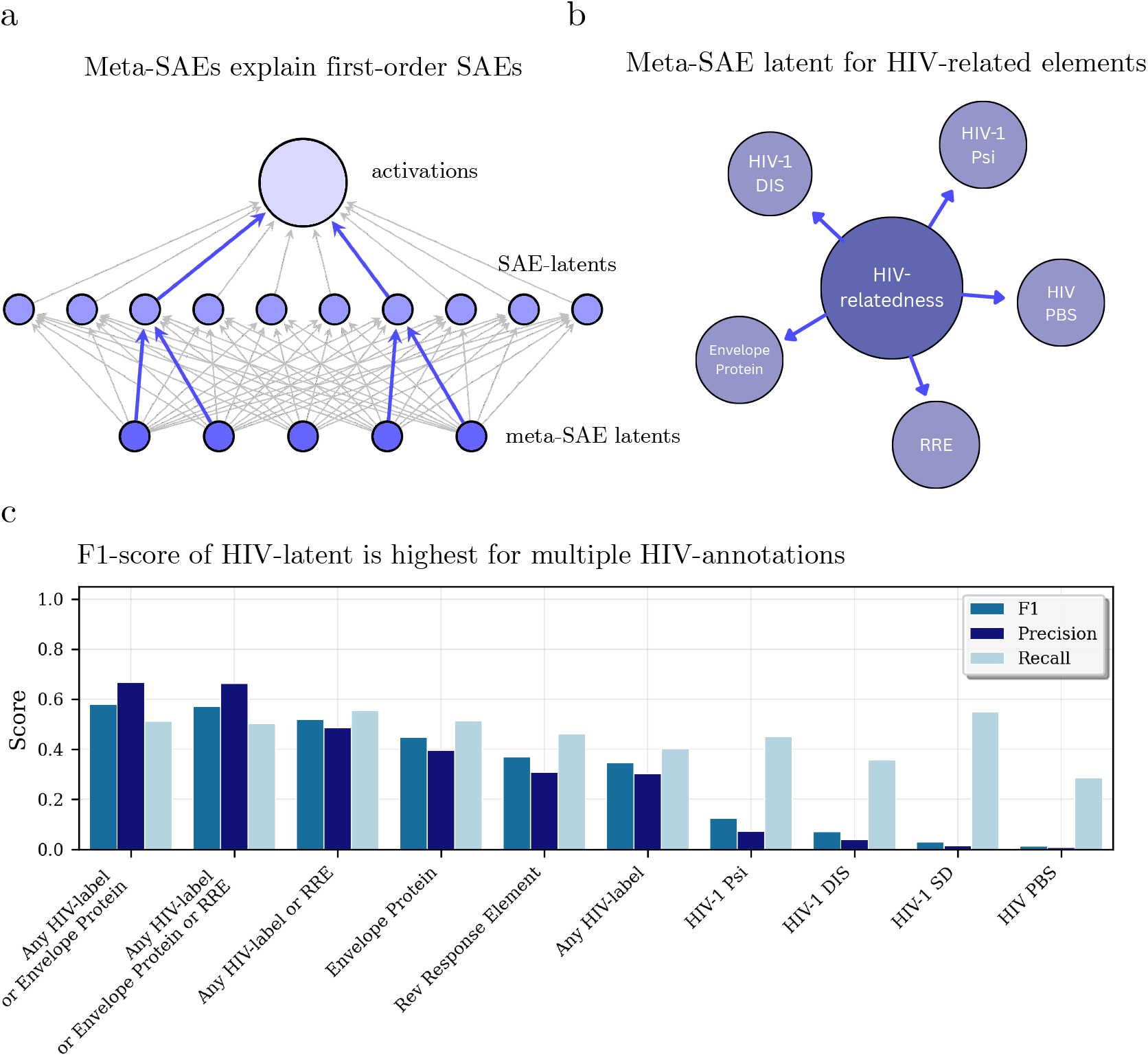
Meta-SAEs identify an abstract concept related to HIV. a) While SAEs are trained to reconstruct model activations, Meta-SAEs are trained to reconstruct the decoder weights of first-order SAE-latents. b) We find a Meta-SAE latent that activates on a cluster of HIV-related first-order SAE-latents. c) We assess this “HIV-latent” by treating its encoder-weights as a probe and measuring the probe F1-score across simple HIV-related annotations and their combinations, finding that the probe best separates a collection of many annotations.

These results indicate that Meta-SAEs recover hierarchical structure within the representations of genomic language models. Whereas the base SAE isolates interpretable, single-annotation features, the Meta-SAE aggregates them into higher-order modules that correspond to coordinated biological processes. For example, latent #107 encapsulates multiple HIV-related sequence elements, RRE, DIS, Psi, PBS, and Env, reflecting the model’s ability to link dispersed regulatory regions. This hierarchical organisation suggests that model features are not independent but form groupings reflecting shared regulatory or structural roles. By quantifying these relationships, Meta-SAEs provide a means to map dependencies between features. In practical terms, this enables more systematic interpretation of genomic language models, revealing how they encode multi-scale biological structure rather than isolated annotations.

### 2.4 Steering toward the A1408G substitution in 16S rRNA with SAE-latents

Having observed that SAE-latents can closely capture known biological annotations, we asked whether individual SAE-latents can causally modulate sequence predictions toward biologically meaningful variants. *Steering* refers to a causal intervention in the model’s activation space, which results in a desired change in the model’s output sequence. Steering involves reconstructing the model’s activations using our SAE, while clamping an SAE-latent to a high value, and then continuing the forward pass with that reconstruction (see Methods 1).

Among features with confident annotations, latent #5066 exhibited a strong association with kanamycin resistance (*F* 1 > 0.6, precision ≈ 0.9). We focused on the well-characterised A1408G substitution in 16S rRNA (Hobbie et al., 2006; Lynch & Puglisi, 2001; Nessar et al., 2011) and evaluated how manipulating #5066 alters the model’s probability assigned to the corresponding “mutation token”. We explore clamping to values that lie in and out of distribution. We define in-distribution steering as setting a latent to values within the empirical range observed on the steered sequence, while out-of-distribution refers to values beyond this range. We perform the steering on layer 10 of the 500M multi-species model, on which the SAE was trained.

At in-distribution steering levels, steering using #5066 at the masked position and also the immediately preceding position increased the A1408G logit and probability, promoting it to Top-1 among candidate tokens. Steering at a distant position (100 tokens before) had no effect, as did steering only on a masked or nearby token (token preceding the mask).

The rank of the mutation token in the non-steered forward pass was already very high at rank 2. Therefore, the change to rank 1 due to some random perturbation may not be an unlikely event. We address this concern by screening other well-annotated latents (F1 ≥ 0.5), which showed that the observed in-distribution effect was selective: only #5066 and #4420 (annotated as SmR, a streptomycin resistance gene) induced a Top-k rank shift, whereas the non-AMR latents did not (Fig. 4d).

**Figure 4:**
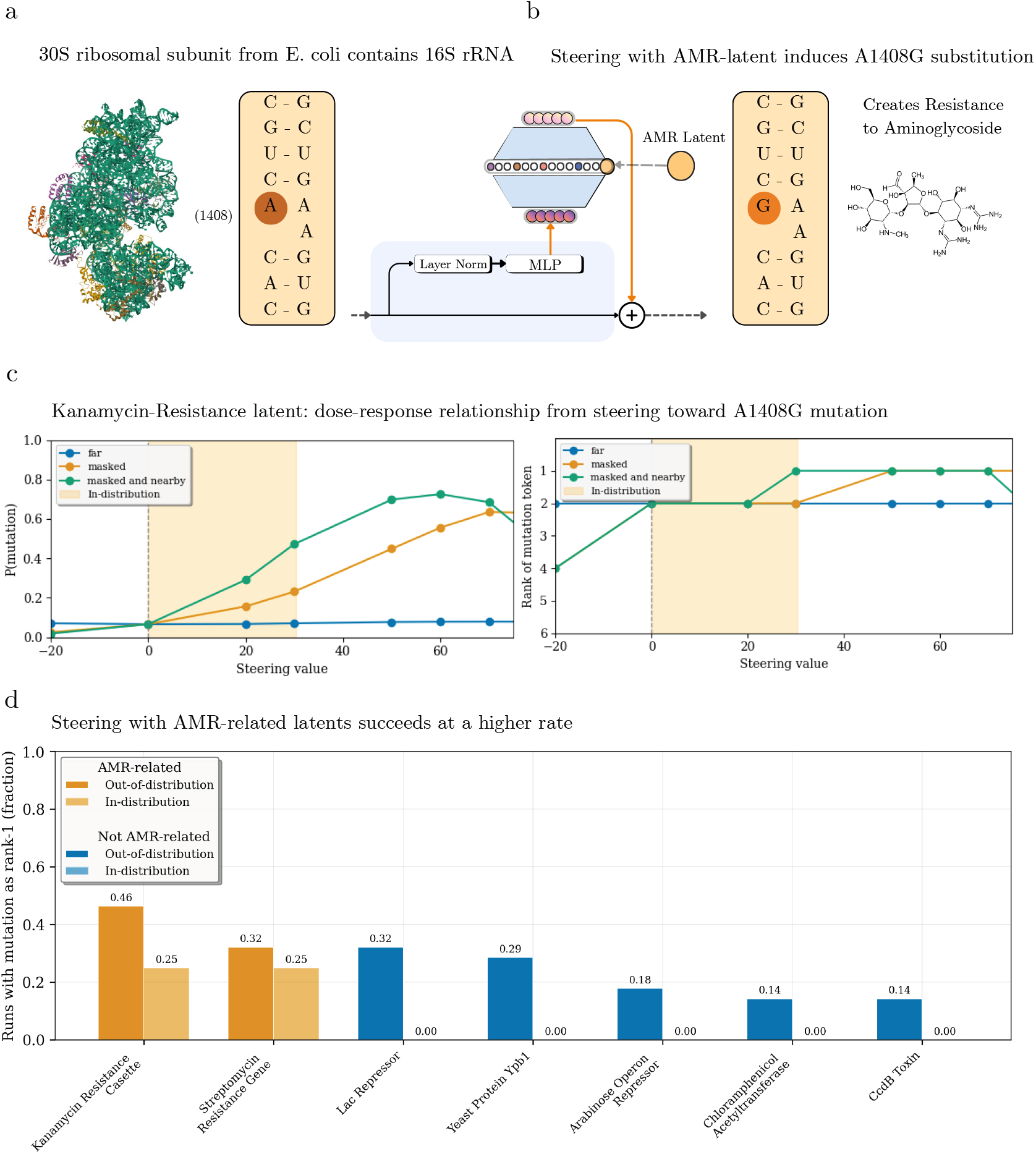
Biologically meaningful steering with a Kanamycin resistance SAE-latent. a) The 30S ribosomal subunit in E. coli (left) contains the 16S rRNA folded onto itself (centre), highlighting nucleotide position 1408, where the A to G substitution confers aminoglycoside resistance. Structure rendered from RCSB Protein Data Bank (PDB ID: 7OE1; (Maksimova et al., 2021)). b) Steering (left) involves modifying the forward pass by reconstructing the MLP outputs using our SAE, with a latent for antimicrobial resistance, clamped to a high value. This induces the A1408G substitution (right), which confers aminoglycoside resistance. Kanamycin A structure (right; image by SSamadi15, Wikimedia Commons, CC BY-SA 4.0). c) Shows that the model shifts probability mass toward the A1408G mutation as we increase the strength of SAE-steering (left) and how the rank of the mutation-token changes (right). Steering on the masked token and nearby tokens leads to a faster increase in probability mass (left) and is necessary for rank changes on in-distribution steering levels (right). d) We find that steering towards the mutation-containing token succeeds at a higher rate for Kanamycin Resistance latent compared to other interpretable SAE-latents. This is especially true if we only steer at in-distribution levels, where the success rate for all other latents becomes zero.

**Figure 5:**
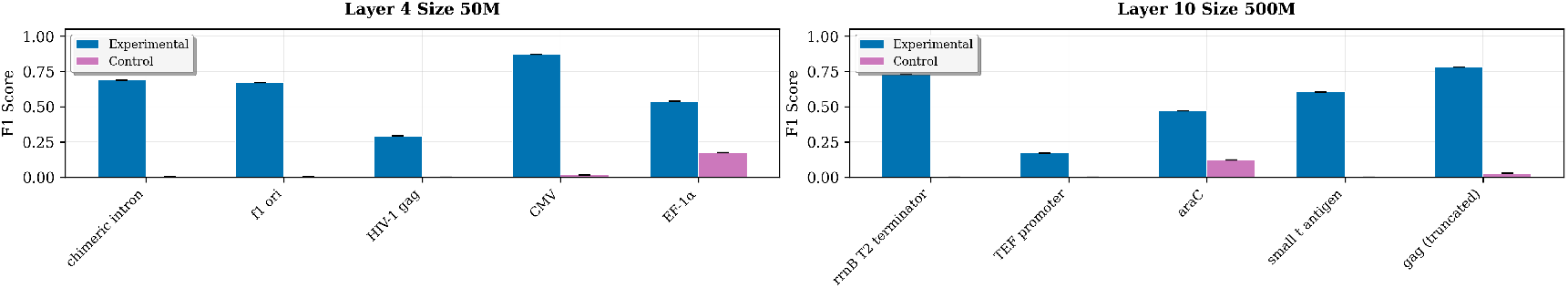
Functional annotations are encoded via pretraining. Comparing linear probe performance across 10 annotations in Layer 4 of the 50M model (left) and Layer 10 of the 500M model (right), consistently shows that the annotations are more linearly separable in the pretrained than in the base model.

We next characterised the dose–response relationship by scaling the steering magnitude beyond in-distribution ranges. The effect size on A1408G increased monotonically up to a breaking point, and sign-reversal (negative steering) suppressed it (Fig. 4a), and for many of the high steering values, steering at either mask or nearby token is sufficient for the rank change. At these higher doses, several non-AMR latents also produced rank shifts (Fig. 4b), indicating that specificity degrades outside the model’s activation distribution. In fact, many of the latents show a dose-dependent effect on the token’s probability similar to that of the antibiotic-resistant latents #5066 and #4420 or an inverted dose-dependent effect where the negative values steer towards greater likelihood.

Together, these results suggest that AMR-annotated features contain directions that can steer predictions toward resistance mutations at realistic activation levels. We note that the SAE has been trained to approximate the activation space of the Nucleotide Transformer. Thus, steering with out-of-distribution values can move our reconstruction out of the space the SAE learned to approximate well and so lead to unexpected effects on the predicted sequence. Furthermore, it has been shown that in artificial neural networks, semantic directions in activation space are not scaling invariant, and so scaling a direction that correlates with some semantic meaning can change the semantic meaning of the vector, especially when moving out of distribution (Black et al., 2022). Together, this might mean that steering works reliably only if we clamp latents to values within the distribution of values present in the sequence, which would be consistent with our observations. Our results also showcase the fragility of steering tasks with respect to the steering method used, especially in biological models where vocabulary size is limited and the underlying meaning of tokens and wider context windows is opaque.

## 3 Conclusion

Our central hypothesis was that genomic language model activations encode a large vocabulary of biological properties in superposition, and that sparse autoencoders can reveal this vocabulary by disentangling their representations. We tested this by training sparse autoencoders on the Nucleotide Transformer and showed that SAE-latents strongly correlate with a diverse array of interpretable biological concepts, uncovering over 60 distinct functional annotations encoded in the model activations. This included viral regulatory elements that we traced to training data contamination, demonstrating that interpretability methods can help diagnose quality issues in large-scale genomic datasets. We then showed that the Nucleotide Transformer encodes concepts at different levels of generality. Using Meta-SAEs, trained on the weights of a base SAE, we showed that the model encodes a more general concept related to HIV. Finally, we showed that the concepts extracted by our SAE can modulate the Nucleotide Transformer’s outputs in biologically plausible ways. In particular, we show that steering with an SAE-latent related to Kanamycin or Streptomycin Resistance causes the model to predict the A1408G substitution in the 16S rRNA, which confers aminoglycoside resistance.

We want to highlight a few important limitations and caveats. First, even our most interpretable SAE-latents have F1-scores markedly below 100%, which may be explained by superposition but could also indicate that the latent directions do not truly correspond to any of the annotations but, instead, some concept that strongly overlaps with it. Second, while our SAE-latents are, in many cases, clearly more interpretable than MLP-neurons, the large majority of SAE-latents do not correlate with a known biological concept. We have not studied these latents in much detail here. Third, while we found contamination of the training data with CMV enhancer sequences, we did not find that contamination for other viral features, leaving them unexplained. Fourth, we do not claim to have found a unique and complete set of features encoded by the model, since annotations are incomplete and SAE features are non-atomic and incomplete as well (Leask et al., 2025). Relatedly, a different plasmid-annotation tool may have led to significantly different latent interpretations in some cases.

Together, these findings establish SAEs as an effective method for analysing representations of genomic language models, with applications in feature discovery, dataset auditing, and tracking representational shifts during finetuning. Future research should apply SAEs to less-annotated regions and develop SAE-steering strategies tailored to the generation of complex sequences. These advances are realistic, given their success in other domains (Templeton et al., 2024; Simon & Zou, 2025), and would position SAEs as a general tool for understanding, auditing, and ultimately designing biological systems.

## Acknowledgements

We want to thank James Black, Joseph Bloom, Nikhil Branson, Sam Brown, Laura Dillon, Clément Dumas, and Julian Minder for helpful discussion and feedback (ordered alphabetically). We would also like to thank funders: AM acknowledges funding by Open Philanthropy, OMC acknowledges funding from a New College Todd-Bird Junior Research Fellowship and MRC Fellowship MR/Y010078/1. He acknowledges consulting fees to Pelago Biosciences, Faculty.ai and MarketCast and is on the scientific advisory board of Evolvere Biosciences. No funder had a role in the research or decision to publish. PJ is supported by funding from the Biotechnology and Biological Sciences Research Council UKRI-BBSRC grant (BB/T008784/1). PJ cofounded Evolvere Biosciences, but the company had no role in this study. FD’s contribution to this work was financially supported by the Hector Fellow Academy. GMM thanks the EPSRC for support via EP/S024093/1.

## 4 Code and Reproducibility

All code developed for SAE training, evaluation, and analysis, along with the trained model weights and scripts to reproduce our main figures, are made publicly available at COMING SOON.

## A Related Work

A central challenge in interpretability research is that neural networks rarely represent input features at the level of individual neurons. Instead, features are encoded as directions in high-dimensional activation space, which do not align with its standard basis directions (neurons), a phenomenon described by the *superposition hypothesis* (Elhage et al., 2022). This allows models to encode more features than neurons, but makes individual activations polysemantic and difficult to interpret directly. One approach to recovering these underlying feature directions is *sparse dictionary learning*, which approximates an activation vector **x** as a sparse, weighted sum of basis directions:

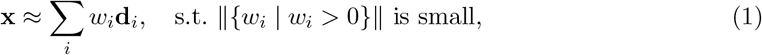

where **d**_*i*_ are dictionary atoms and *w*_*i*_ are sparse coefficients.

Sparse autoencoders (SAEs) are one implementation of this idea. Formally, SAEs consist of an encoder *f*_*θ*_: ℝ ^*n*^ → ℝ ^*m*^ and a decoder *g*_*ϕ*_: ℝ ^*m*^ → ℝ ^*n*^ trained to minimize the reconstruction error

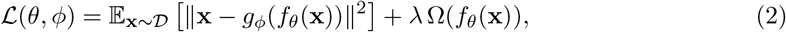

where Ω is a sparsity-inducing regularizer and *λ* controls its strength. By learning an overcomplete dictionary under a sparsity constraint, SAEs are intended to disentangle superimposed features and yield more interpretable latent units.

We compare SAEs to three alternative approaches to interpreting the gLMs’ latent representations. First, researchers have applied *unsupervised clustering* of sequence embeddings with UMAP (Benegas et al., 2023; Chen et al., 2024) or t-SNE (Nguyen et al., 2023). Second, *attention weight analysis*, such as in Chen et al. (2024), provides insights into long-range dependencies between sequence elements but, unlike SAEs, does not directly isolate disentangled features in the model’s representation space. Third, many papers have trained classifiers (*probes*) on hidden representations of the model to test if the model representations encode various functional genetic elements (Benegas et al., 2023; Dalla-Torre et al., 2025; Tang et al., 2024; Hwang et al., 2024). Unlike SAEs, probing classifiers require supervised labels, making them well-suited for testing pre-defined hypotheses about what features a model encodes. However, this reliance on labeled data limits their usefulness for discovering novel, unexpected representations - a key advantage of SAEs.

By identifying explicit directions in the representation space, SAEs enable interventions such as *model steering or unlearning* specific encoded features. However, recent work in the domain of natural language suggests that alternatives outperform SAE-based methods for both purposes: Wu et al. (2025) suggest that alternatives like prompting techniques are more effective for steering and Farrell et al. (2024) suggests that Representation Misdirection for Unlearning (Li et al., 2024) provides several advantages for unlearning.

We focused this section on gLM interpretability techniques as applied to model representations. This does not cover other interpretability techniques, e.g. methods for characterizing the relative importance of various input parts to the model’s output. For a more comprehensive review of interpretability methods for gLMs, see Benegas et al. (2025).

Most models previously interpreted with SAEs were Transformers trained on large corpora of text (Yun et al., 2023; Cunningham et al., 2023). Recently, this approach has been extended to protein Language Models, ESM-2 (Simon & Zou, 2025), and even the genomic language model EVO 2 (Brixi et al., 2025). In contrast to (Brixi et al., 2025), we apply this method to a standard Transformer-based gLM that is pretrained with a masked language modelling objective: the Nucleotide Transformer from Dalla-Torre et al. (2025).

A key challenge in understanding the representations learned by Sparse Autoencoders (SAEs) is interpreting the latent activations in a meaningful way. Previous work primarily relied on manual inspection of the most activating inputs to identify unifying concepts behind each latent dimension or used large language models (LLMs) to automate this process (Bricken et al., 2023; Gao et al., 2024). While this approach has been effective in domains such as natural language and vision, it is less suitable for biological sequence data, where meaningful features often correspond to functional or regulatory elements that are not trivially observable from raw sequence patterns. To address this limitation, we follow Simon & Zou (2025) in using automatically annotated biological features as an intermediate step before interpreting SAE-latents. These annotations allow us to quantitatively measure the association of an SAE latent and a concept.

In designing the SAE architecture, the choice of activation function plays a crucial role in the effectiveness of SAEs at extracting interpretable features. Early SAE implementations primarily used ReLU activations (Cunningham et al., 2023; Bricken et al., 2023), which enforce sparsity indirectly by thresholding negative activations at zero. To promote sparsity while avoiding “activation shrinkage”, the Jump-ReLU activation function was proposed by Rajamanoharan et al. (2024). An alternative line of work has explored activation functions that fix the number of active latents for a set of inputs. Gao et al. (2024) introduced Top-K activation functions, setting all but the K largest activations for a particular input to zero. More recent work by Bussmann et al. (2024) has modified this approach by allowing the top *k × n* latents to be active across a sample of *n* inputs, enabling more flexibility in the number of dictionary elements used to reconstruct a single input. We choose the BatchTopK activation function as it is widely regarded as being more effective than TopK and easier to train than JumpReLU SAEs.

As introduced by Bussmann et al. (2024), BatchTopK Sparse Autoencoder (BatchTopK SAE) is a variant of the TopK sparse autoencoder that enforces sparsity at the *batch level* rather than per sample. Given an input batch *X* ∈ ℝ^*n×d*^ consisting of *n* samples, the encoder produces activations

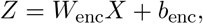

where *W*_enc_ ∈ ℝ ^*d×m*^ and *b*_enc_ ∈ ℝ ^*m*^. The BatchTopK operator retains the top *n × k* activation values across the entire batch, setting all others to zero:

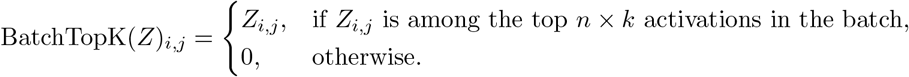

The decoder reconstructs the batch as

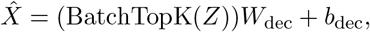

where *W*_dec_ ∈ ℝ ^*m×d*^ and *b*_dec_ ∈ ℝ ^*d*^. The training objective minimizes the reconstruction loss with an auxiliary regularization term to prevent dead latents:

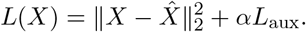

During training, the encoder computes activations *Z*, after which the BatchTopK operator selects the globally top activations across all samples in the batch. This allows individual samples to activate a variable number of latents, improving reconstruction quality while maintaining a fixed *average sparsity*. During inference, the batch dependency is removed by estimating a global activation threshold

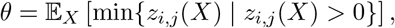

and replacing the BatchTopK operator with a JumpReLU-style activation:

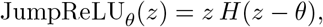

where *H*(·) is the Heaviside step function. This yields a fixed thresholding rule that preserves the sparsity structure learned during training.

A *Meta-SAE* (Leask et al., 2025) extends the sparse autoencoder framework by learning higher-level structure among features discovered by a base SAE. Instead of being trained on raw activations, it is trained on the decoder weights of a previously trained SAE. Let 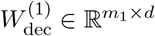 be the decoder of the base SAE, where each row 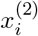 represents a learned feature in the original activation space ℝ ^*d*^. Stacking these feature vectors gives the dataset for the Meta-SAE:

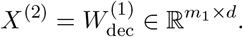

The Meta-SAE defines an encoder-decoder pair:

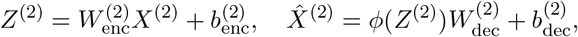

where 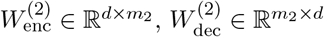, and *ϕ*(·) is a sparsifying activation such as JumpReLU or TopK. The training objective minimises the reconstruction error of the base decoder weights:

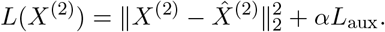

Each base feature (row of 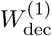) is thus reconstructed as a sparse linear combination of meta-features (rows of 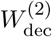):

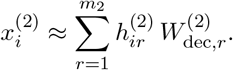

where 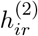 are the sparse activations produced by the Meta-SAE encoder.

## B Methods

### B.1 Experimental Setup

#### Choice of Nucleotide Transformer

We chose to focus on the Nucleotide Transformer (Dalla-Torre et al., 2025). One advantage of this model family is that it spans multiple scales, including a relatively small 50m parameter model, with relatively low-dimensional hidden states. This is advantageous because the computational cost of training SAEs grows linearly with the hidden-state size of the model (all else equal). A second reason for the Nucleotide Transformer family is that it has demonstrated strong benchmark performance, indicating that it has successfully encoded a variety of input properties, making it particularly promising for interpretability research.

#### Choice of Dataset

The most obvious dataset for training a sparse autoencoder on the Nucleotide Transformer is the model’s pretraining data, a dataset of 850 genomes, spanning multiple domains of life. However, functional annotations for these genomes are scarce, which results in most SAE-latents activating on regions without known function, such as hypothetical proteins. To avoid this problem, we chose to focus on sequences that can be densely and confidently annotated: plasmids. We downloaded a dataset of plasmids from AddGene, an online repository for genetically engineered plasmids (Kamens, 2015). Our dataset consisted of 93,306 plasmids for training and about 15,551 for validation and testing.

#### Creating Annotated Evaluation Sets

To functionally annotate plasmid sequences, we used pLannotate (McGuffie & Barrick, 2021), a specialised tool for automated annotation of plasmid features. Our annotation pipeline consisted of the following steps:

1. Dataset preparation: We randomly selected three non-overlapping sets of 1,000 plasmid sequences each from a larger validation dataset of Addgene plasmids. To ensure reproducibility, we used a fixed random seed (42) for the selection process.
2. Annotation procedure: Each plasmid sequence was processed using pLannotate’s annotate function with parameters set to include detailed annotations and treating sequences as linear. This procedure identified functional genetic elements including promoters, resistance markers, replication origins, and other standard plasmid features.
3. Annotation output: For each plasmid, the annotation process generated a comprehensive table of hits containing feature coordinates, feature types, and confidence scores (e-values)
4. Data organization: We compiled annotation results for each set into separate CSV files, preserving sequence identifiers for cross-referencing. These annotation files served as the foundation for subsequent analyses of feature representation in the Nucleotide Transformer activations.

All annotation processing was performed using Python 3.8 with pLannotate v1.2.0, pandas v1.3.5, and BioPython v1.79. We performed almost all our training in Google Colabs, using either a single A100 (for SAE training) or a single L4 GPU (for linear probe training).

### B.2 Training Sparse Autoencoders

For each plasmid sequence, we extracted MLP outputs at layer 1, 4, 7, 10 and 12 (final) of the 50m Nucleotide Transformer and at layer 10 of the 500m Nucleotide Transformer. We downloaded the models from HuggingFace.

As mentioned before, we used BatchTopK SAEs, with an expansion factor of 8. We trained each SAE for 100,000 steps with a batch size of 4096 and a maximum sequence length of 512 tokens. We used the Adam optimiser with learning rates set to 3e-4 and linear warmup over the first 10% of steps and linear decay over the last 20%. We set K-values to 16 or 32. We normalised activations by 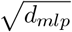. Our goal was to demonstrate that SAEs can extract biological concepts from the Nucleotide Transformer, rather than finding the “best” SAE for this model. We used the hyperparameters from Bussmann et al. (2024).

Following the implementation of (Bussmann et al., 2024), we initialised the encoder as the transpose of the decoder weights. We maintained unit-norm decoder columns throughout training, while also performing gradient-clipping and removing gradients parallel to decoder columns before each update.

### B.3 Evaluating Sparse Autoencoders

To screen promising SAEs, we initially relied on three proxy metrics: (1) normalised MSE, measuring reconstruction fidelity, (2) L1-loss, measuring sparsity, and (3) the fraction of dead latents, activating less than once per 10 million tokens. We detail these below for each of our SAEs.

To determine if our SAE-latents had successfully captured known biological concepts, we had to correlate activation of individual SAE-latents with the annotations of activating input tokens. For our analysis, a token (a 6-mer) was considered annotated if it shared at least one nucleotide with a region identified by pLannotate.

#### Inspection of Individual Latents

To understand the function of individual SAE-latents, we examined the tokens that elicited the strongest activations for each latent. For each highly activating token, we recorded its annotation (if any), its surrounding sequence context, the annotation of that context, and a confidence score (e-value) associated with the annotation. This allowed us to qualitatively assess whether a latent was responding to specific biological features.

#### Measuring Monosemanticity

A key aspect of evaluating the learned SAE-latents is to quantify their monosemanticity – the degree to which each latent corresponds to a single, well-defined concept. While perfectly monosemantic neurons, which activate exclusively for a specific property, are rarely observed in practice, we aim to measure how closely our latents approach this ideal. For instance, a highly monosemantic latent for “promoter” would ideally activate strongly and consistently for promoter sequences and remain inactive for other types of sequences. To quantify monosemanticity, we adopted the standard F1-score metric, *in contrast to* Simon & Zou (2025) who used “Domain F1-score”. We didn’t find persuasive reasons to adopt Domain F1-scores, and since we found them to be generally much higher, we were concerned that they might inflate how strongly our latents are associated with respective annotations.

#### Domain Recall vs. Standard Recall

The primary difference between Domain F1-score and the standard F1-score lies in the calculation of recall. Standard recall measures the proportion of true positive tokens (tokens with a specific annotation) that a classifier correctly identifies. In contrast, Domain Recall focuses on “true domains” – contiguous regions of multiple tokens that are all annotated with the same functional element. Domain Recall calculates the fraction of these true domains in which a given latent activates on at least one constituent token. This modification significantly increases the measured recall, and consequently the F1-score, of our latents. The difference between these two recall metrics and their impact on the F1-score for our SAE-latents is visually illustrated in Figure 2. As expected, we generally observe lower values for the standard metrics. Notably, latents associated with puromycin resistance, envelope proteins, and streptomycin resistance maintain relatively high F1 values even under the standard metric.

#### F1-score with Activation Threshold

We swept thresholds over the observed activation range and selected the value that maximised validation F1. Importantly, the final F1-scores reported in our study are computed on two non-overlapping hold-out test sets to avoid overfitting.

### B.4 Concept Validation Through Probing

For each concept, we train 30 logistic regressions on both the MLP outputs of the Nucleotide Transformer and a randomly initialised version. We train for 50 epochs, with early stopping after no improvement of at least 1e-5 in F1-score after 20 epochs. We use a batch size of 128. Detailed results are shown in Figure 5.

### B.5 Training Meta-SAEs

We train Meta-SAEs using three sparsity mechanisms: BatchTopK, JumpReLU, and classical Lasso-based dictionary learning (Mairal et al., 2009). All SAEs are trained with a batch size of 512, a learning rate of 2 *×* 10^−4^, and for 5000 optimisation steps, using a linear warmup of the *ℓ*_1_ penalty over the first 1000 steps. For JumpReLU models, we set the bandwidth to 0.01 and the initial threshold to 0.001. Encoders are initialised as the transpose of their corresponding decoders, and all weights use Kaiming Uniform initialisation (He et al., 2015). Across configurations, JumpReLU SAEs achieve slightly better reconstruction scores than BatchTopK or Lasso-based dictionary learning. Increasing the latent dimensionality generally improves reconstruction performance, with 128 latent units providing a good balance between compactness and accuracy.

### B.6 SAE-based Steering

We assess whether sparse autoencoder (SAE) latents can be used to steer a masked language model (MLM) by directly manipulating latent activations at a chosen layer and reinjecting a residual-preserving reconstruction into the forward pass to obtain steered logits.

#### Model and layer

Steering is applied to the Nucleotide Transformer (NT) at a single unique layer during masked language modelling.

#### Overview of steering procedure

Given an input sequence *s* and a layer *j*:

1. **Capture baseline activations:** Run a forward pass and hook the MLP output at layer *j* to obtain per-position activations **a** ∈ ℝ ^*N×d*^, and record control logits **z**^ctrl^ ∈ ℝ ^*N×V*^.
2. **Decompose with SAE:** Pass **a** through *SAE* to obtain

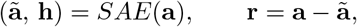

where **h** are the hidden states of the SAE, **ã** is the reconstruction of the activations and **r** is the reconstruction residual.
3. **Manipulate latents:** Let *I*_max_ be indices of latents to amplify and *I*_0_ indices to set to zero. Optionally restrict to a set of token positions *P* ⊆ {1,…, *N*}. For *i* ∈ *I*_0_ set *h*_*n,i*_ ← 0; for *i* ∈ *I*_max_ set

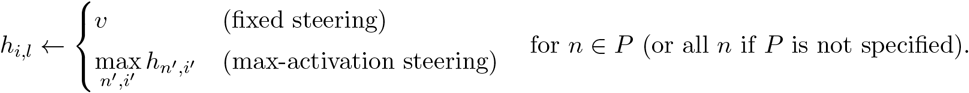
4. **Residual-preserving reconstruction:** Decode the edited latents and add back the residual:

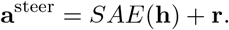
5. **Steered forward pass:** Inject **a**^steer^ at layer *j* via a forward hook and complete the pass to obtain steered logits **z**^steer^.

#### Token- and vocabulary-level effects

We compute the logit difference

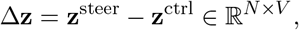

its mean over sequence positions 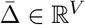, and report the top-*k* increased and decreased vocabulary items by entries of 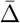.

#### Latent manipulation policies

Two steering-value policies are supported: (1) *fixed* (*v* ∈ ℝ_*>*0_), and (2) *max-activation*, which sets selected latents to the maximum latent value observed in the current example. Steering can be applied globally or only at designated positions *P* (e.g., masked or motif-aligned tokens). We find that steering on the entire sequence leads to poor results.

#### Implementation details

Hooks intercept and substitute the output of the layer-*j* MLP projection (output.dense). Shapes are matched per batch/sequence; hooks are cleared after each pass. Returned diagnostics include **z**^ctrl^, **z**^steer^, lists of top increased/decreased tokens, and 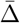.

##### Algorithm 1

SAE Latent Steering with Residual-Preserving Injection

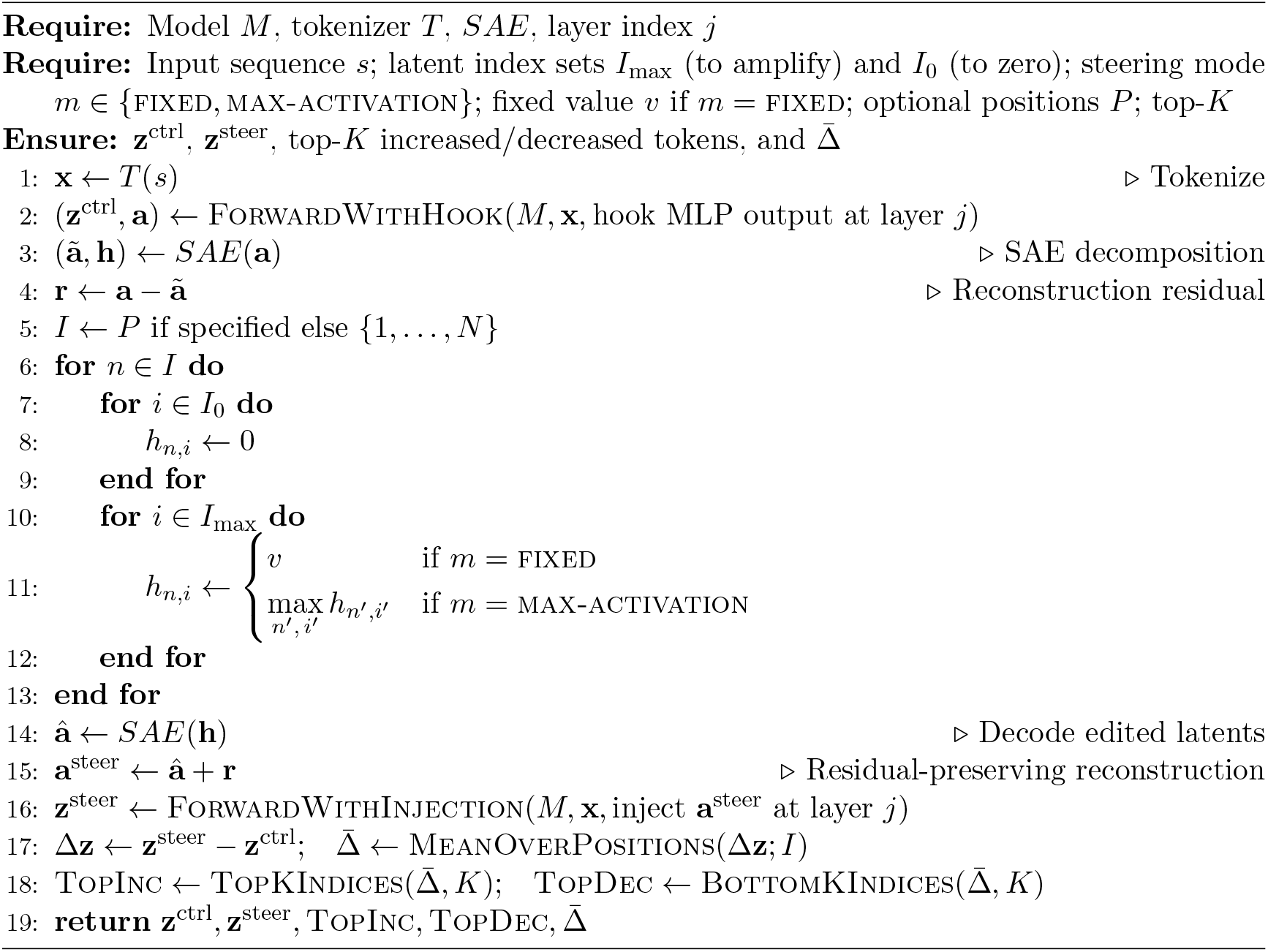

#### Outputs and reporting

We report (i) control vs. steered logits, (ii) 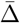 over the vocabulary, and (iii) the top-*K* most increased/decreased tokens by 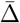. When steering is position-restricted, summaries are computed over *i* ∈ *P*; otherwise over all positions.

## C Supplementary Information

### C.1 Characterising the Sequence Features Underlying Probe Performance

To understand what sequence features the CMV enhancer probe relies on, we tested two hypotheses: that it recognises specific regulatory motifs, or that it responds to simpler sequence statistics.

#### Motif ablation reveals minimal contribution of known binding sites

We systematically ablated four known motifs in CMV enhancers—E-box sequences, CREB/ATF sites, NF-*κ*B binding sites, and CCAAT boxes—replacing each with random sequences of identical length. Only E-box ablation significantly reduced probe probability compared to both original and control sequences with random ablations (*p* ≤ 0.0001, Wilcoxon signed rank test). The other three motifs showed no significant effect, suggesting the probe does not primarily rely on these canonical regulatory elements.

#### K-mer statistics explain probe performance

We next tested whether simpler sequence features could account for probe behavior by generating decoy sequences that preserve k-mer frequencies while disrupting higher-order structure. For each *k* from 1 to 5, we shuffled sequences to maintain exact *k*-mer counts while randomizing their positions. Figure 6B shows that probe performance increased monotonically with *k*: preserving only nucleotide (*k* = 1) or dinucleotide (*k* = 2) frequencies yielded poor performance, while *k* = 5 recovered 85% of the original probe probability. This 5-mer dependency explains why longer motifs (6–12 bp) contribute minimally—the probe captures local sequence patterns rather than extended regulatory elements.

**Figure 6:**
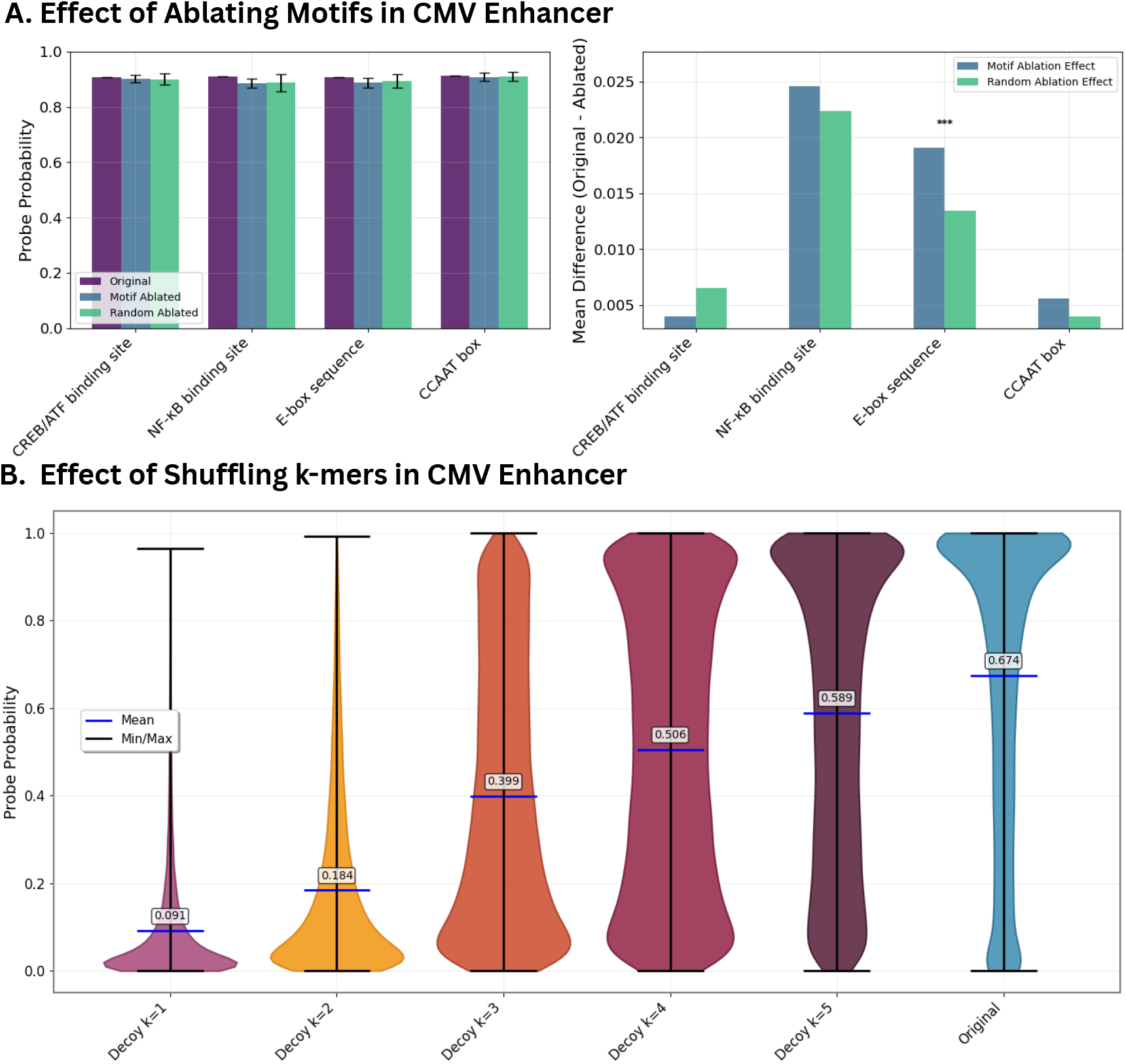
The Effect of Disrupting the Sequences on CMV Enhancer Prediction. A) Shows the effect of ablating motifs from CMV enhancer sequences on the probe probability assigned to these sequences. As baselines, we compare to the original sequence and a sequence with a random ablation of the same length. B) Compares the average probe probability for CMV enhancer sequences to decoys, shuffled sequences that preserve only the distribution of k-mers, not their order.

## Footnotes

1 Specifically, we train on the MLP outputs.

2 In the large-sample independence approximation, if *f* is the frequency (probability) that a token carries annotation *A* and a latent fires independently with probability *r*, then treating probabilities as expected token fractions gives Pr[TP] = *fr*, Pr[FP] = (1 − *f*)*r*, and Pr[FN] = *f* (1 − *r*). Hence precision *P* = *f* and recall *R* = *r*, and the expected F1 (harmonic mean) is 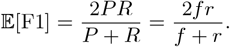 Maximising this over *r* ∈ [0, 1] yields *r* = 1 and gives the upper bound 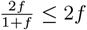. Because all of our annotations have 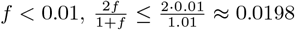, so the random-baseline expected F1 is at most ≈ 1.98% (reported above conservatively as 2.0%).

